# Spatial and epidemiologic features of dengue in Sabah, Malaysia

**DOI:** 10.1101/657031

**Authors:** Amanda Murphy, Giri Shan Rajahram, Jenarun Jilip, Marilyn Maluda, Timothy William, Wenbiao Hu, Simon Reid, Gregor J. Devine, Francesca D. Frentiu

**Affiliations:** Mosquito Control Laboratory, QIMR Berghofer Medical Research Institute, Brisbane, Australia; School of Biomedical Sciences, and Institute for Health and Biomedical Innovation, Queensland University of Technology, Brisbane, Australia; Queen Elizabeth Hospital, Ministry of Health Malaysia, Kota Kinabalu, Malaysia; Infectious Disease Society of Kota Kinabalu-Menzies School of Health Research Clinical Research Unit, Kota Kinabalu, Malaysia; Sabah Department of Health, Ministry of Health Malaysia, Kota Kinabalu, Malaysia; Gleneagles Kota Kinabalu Hospital Sabah, Kota Kinabalu, Malaysia; School of Public Health and Social Work, Queensland University of Technology, Brisbane, Australia; School of Public Health, University of Queensland, Brisbane, Australia

**Keywords:** dengue, rural, Sabah, Aedes albopictus, Borneo, South East Asia

## Abstract

In South East Asia, dengue epidemics have increased in size and geographical distribution in recent years. Most studies investigating dengue transmission and control have had an urban focus, while less consideration is currently given to rural settings, or where urban and rural areas overlap. We examined the spatiotemporal distribution and epidemiological characteristics of reported dengue cases in the predominantly rural state of Sabah, in Malaysian Borneo – an area where sylvatic and urban circulation of pathogens are known to intersect. We found that annual dengue incidence rates were spatially variable over the 7-year study period from 2010-2016 (state-wide mean annual incidence of 21 cases/100,000 people; range 5-42/100,000), but were highest in rural localities in the western districts of the state (Kuala Penyu, Nabawan, Tenom and Kota Marudu). The eastern districts exhibited lower overall dengue rates; however, we noted a concentration of severe (haemorrhagic) dengue cases (44%) in Sandakan and Tawau districts. Dengue incidence was slightly higher for males than females, and was significantly higher for both genders aged between 10 and 29 years (24/100,000; p=0.029). The largest ever recorded outbreaks occurred during 2015-2016, with the vector *Aedes albopictus* found to be most prevalent in both urban and rural households (House Index of 64%), compared with *Ae. Aegypti* (15%). These findings suggest that dengue outbreaks in Sabah are driven by the sporadic expansion of dengue virus in both urban and rural settings. This may require tailoring of preventative strategies to suit different transmission ecologies across Sabah. Further studies to better understand the drivers of dengue in Sabah may aid dengue control efforts in Malaysia, and more broadly in South East Asia.

**Author summary:** In order to combat the rising regional incidence of dengue in South East Asia, the drivers of transmission must be better characterised across different environmental settings. We conducted the first retrospective analysis of dengue epidemiology in the predominantly rural state of Sabah, Malaysia, where both urban and sylvatic transmission cycles exist. Human notification data over a 7-year period were reviewed and spatiotemporal and demographic risk factors identified. We found:

1. Urban habitats and population density are not the only determinants mediating the spread of epidemic dengue in Sabah. Case from both urban and rural localities contributed equally to dengue outbreaks.
2. Human demographic risk factors included being aged between 10 and 29 years, and being male.
3. High incidence areas for dengue do not predict the occurrence of severe dengue. Severe dengue was largely localised to lower incidence districts in the east of the state.
4. The sole presence of *Aedes albopictus* in and around the majority of urban and rural case households suggests that this vector may play a major role in facilitating outbreaks.

A complex interplay of risk factors likely mediates dengue transmission in Sabah, influenced by both regional climate trends and localised human and ecological influences. This study emphasises that the increasing spread of dengue in urban South East Asia is also mirrored in more rural areas, and suggests a need for control strategies that address both urban and rural dengue risk.

## Introduction

Dengue is the most rapidly spreading vector-borne disease in the world, and the most prevalent arboviral disease of humans (1). Now endemic in more than 100 countries, the disease causes an enormous burden on communities and health care systems in tropical and sub-tropical regions (2). The causative agent of dengue is dengue virus (DENV), transmitted between humans by *Aedes* mosquitoes across a range of domestic and sylvatic environments. Urban expansion, human migration, travel and trade have facilitated an increasing number of infections, primarily in Asia, Africa and the Americas (3, 4). These areas experience up to 70% of the estimated 390 million annual dengue infections worldwide (1, 4). Explosive outbreaks have become common in recent decades, and both classical and severe (haemorrhagic) forms of dengue now occur in previously unaffected countries (1, 3, 5). South East Asia has one of the highest burdens of dengue, following marked increases in the number, severity and geographical distribution of dengue epidemics since the 1950s. During this dramatic expansion, the four virus serotypes (DENV 1-4) have become well-established and commonly co-circulate within the region (6).

In Malaysia, dengue has been considered a major public health problem since 1973 (7), with regular epidemics resulting in significant morbidity and economic burden (8, 9). The majority of reported cases are concentrated in the large, urban cities of Kuala Lumpur and Penang, which are located on the Malaysian peninsula. The circulation of all DENV serotypes has been documented across the country, as well as the presence of unique sylvatic strains (10–12). As with many South East Asian countries, the characterisation and control of transmission in Malaysia is primarily focused on highly populated urban areas (13). The majority of spatial and eco-epidemiological studies to date have therefore focused on peninsular Malaysia, and relatively few studies have explored the factors driving transmission in rural parts of the country.

The Malaysian states of Sabah and Sarawak, located on the island of Borneo, report lower incidence rates than mainland Malaysia (14) and patterns of transmission in these states are not well characterised. The island possesses rapidly developing urban areas in close proximity to disturbed forest environments, with potential risk of spill-over of sylvatic pathogens to human populations (15). Sabah state, positioned on the northern tip of Borneo, reports the highest incidence of the sylvatic malaria parasite *Plasmodium knowlesi*, with transmission risk linked to deforestation (16, 17). The emergence of other zoonotic pathogens has also been documented in Sabah (18, 19), including Zika virus in 2015 (20). Given the marked environmental change occurring in Sabah, and the increase in dengue cases noted in recent years (12, 14), it is essential from a public health perspective to understand current transmission patterns and their drivers. This study examined the epidemiology of dengue in the state of Sabah, in Malaysian Borneo, between 2010 and 2016. We aimed to document recent spatial and temporal trends of dengue disease, and to identify some potential risk factors driving DENV transmission and spread in this understudied region of the country.

## Methods

### Ethics statement

This study was approved by the Medical Research and Ethics Committee (MREC), Ministry of Health Malaysia; and the Human Research Ethics Committee (HREC) of the QIMR Berghofer Medical Research Institute, Brisbane, Australia. All human case data analysed were anonymized.

### Study site

The Malaysian state of Sabah lies at the most north-eastern tip of the island of Borneo. It borders the Malaysian state of Sarawak and the Indonesian province of Kalimantan (Fig. 1). The climate is tropical, with high humidity and rainfall throughout the year. Sabah has a geographical area of 73,904 km^2^ and is divided into 25 districts (21). The state’s population density is second lowest in the country (44 people/km^2^), after Sarawak (20 people/km^2^), and Sabah also has one of the lowest overall proportions of urban population (54%) in the country (14). Within Sabah, Kota Kinabalu district has the highest population density (1,397 people/km^2^), where the capital city of the same name is located.

**Fig 1.**
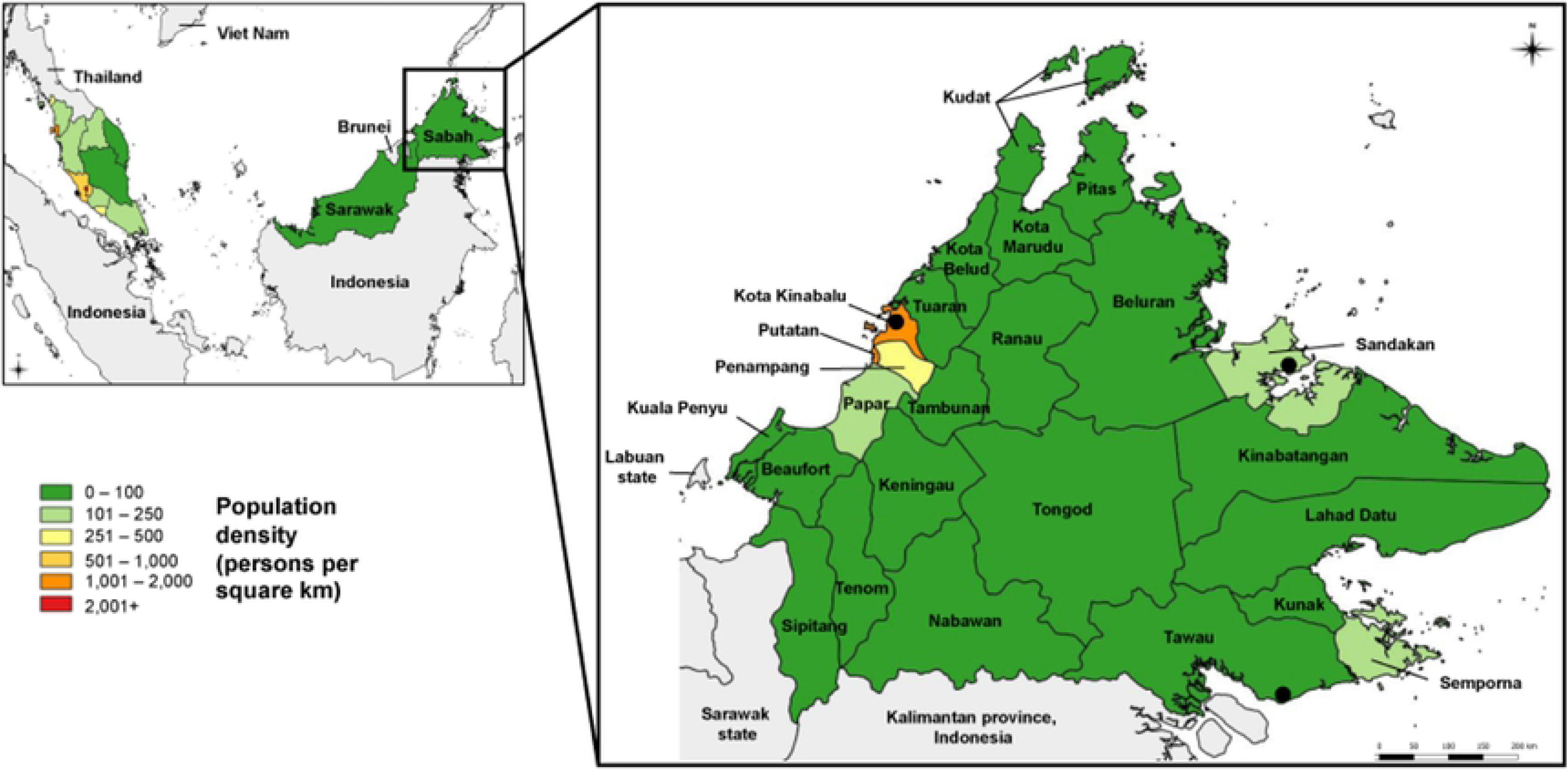
Map of Malaysia and Sabah state. Peninsula Malaysia and Malaysian Borneo are shown, along with the 13 Malaysian states and 2 territories. States are coloured according to their population density, expressed as number of people per square km. The island of Borneo includes the Malaysian states Sabah and Sarawak, and is also shared by the country of Brunei and the Indonesian province of Kalimantan. Inset: Sabah state, showing its 25 districts. The three largest cities in the state are indicated by black circles: the capital city Kota Kinabalu, Sandakan and Tawau.

Historically, Sabah was almost entirely covered by primary rainforest and still has the second-highest proportion of forested areas in the country (60%) after Sarawak (64%). It also has high rates of forest loss, with monocultures of rubber and palm plantations now estimated to cover 36-56% of the land area (15, 21). Sabah has the second-highest proportion of Indigenous people in the country (25%) after Sarawak (30%), as well as the highest proportion of non-Malaysian residents (25%) of all the Malaysian states (14).

### Epidemiological data

State-wide data from monthly notified cases of dengue between the years 2010 and 2016 were obtained from the Sabah State Department of Health (Jabatan Kesihatan Negeri Sabah), Malaysian Ministry of Health. Prior to 2010, detailed data were not available in disaggregated and electronic format. Variables analysed included age, sex, district and locality of each case residence (based on home address), disease severity and outcome (survival or death), and diagnostic tests performed (IgG, IgM and/or NS1). In our dataset, cases from 2011-2016 were designated as residing in either urban or rural localities (the smallest residential geographical unit) by the Sabah Ministry of Health (MoH). MoH designation of locality status is based on the Malaysian Department of Statistics definitions, where urban localities are gazetted census areas with 10,000 people or more, with ≥60% of the working population (≥15 years) engaged in non-agricultural activities (14). Population and demographic data were obtained from the Malaysian Department of Statistics, for the year 2010. Incidence rates were calculated using population projections for each year, based on census data from the year 2010, along with annual growth rate projections as per the published growth rate in Sabah (22).

During the study period, clinical cases were identified using World Health Organization (WHO) guidelines using clinical symptoms and/or positive NS1 or serology (presence of IgM or IgG) (23). From 2014 onwards, Malaysian national notification guidelines were modified, in line with WHO advice, to require a positive laboratory diagnostic test (either NS1 and/or IgM/IgG serology) in addition to the presence of clinical symptoms, and case notification within 24 hours of diagnosis (24, 25). Therefore, the majority of cases prior to 2014 were clinically diagnosed (with 30-50% per year confirmed by laboratory tests in our dataset), while cases from 2014-2016 were 100% laboratory confirmed.

### Entomological data

Entomological surveillance data (number of larvae, mosquito species identified) were generated from active surveillance of potential aquatic habitats, primarily water-holding containers, in and around 719 case residences inspected during the 2015-16 outbreaks. Of these, 255 (36%) of residences were in a locality designated as urban by local public health authorities, 437 (61%) were considered rural, and 27 (4%) had no rural or urban designation recorded. Where mosquito larvae were found in or around a case household, samples were taken to local public health laboratories for species identification. The presence or absence of one or more species per household was recorded, and the House Index (HI) was calculated as the proportion of houses infested with larvae and/or pupae (26). HI was also calculated for each mosquito species present in larvae-positive households.

### Data analysis

We assessed seasonal characteristics of the temporal distribution of cases using a seasonal trend decomposition procedure in SPSS software. The procedure is based on the Census Method I, otherwise known as the ratio-to-moving-average method where time series data are separated into a seasonal component, a combined trend and cycle component, and an “error” or irregular component (27). The seasonal component is then isolated from the overall and irregular trends through a multiplicative model. Seasonal decomposition analysis was applied to monthly dengue case numbers across the 7-year period to examine the seasonal trends of case notifications across Sabah.

Annual and cumulative incidence of dengue was calculated using the number of notifications per month and Sabah population estimates based on the 2010 Malaysian census. Incidence rates were standardized for age and sex using census data and plotted for each of the 25 administrative districts. Ages of cases were grouped into four categories to broadly separate young children from older children and adults (0-9, 10-29, 30-49 and ≥50). The statistical significance of observed differences between means was determined using the Kruskal Wallis test. SPSS Statistics software (SPSS, IBM New York USA; version 23) was used for data analyses, with statistical significance set at p<0.05. Spatial maps of Malaysia and Sabah dengue cases and incidence were created using ArcGIS (Esri Redlands USA; version 10.5.1).

We assessed overall and annual trends of rural versus urban cases at the state-wide level for a 6-year period where locality status was available (2011-2016). This included a total of 9,791 cases. Of these, 756 (7.7%) cases were missing a designated locality status (rural or urban). We classified these cases with no locality status as having ‘unspecified’ localities, and excluded these from rural-urban incidence calculations. For the remaining 9,035 cases, we calculated the total proportions and incidence rates for urban and rural cases, using population projections calculated from state-wide rural-urban population data published in 2010 (22). At district level, we calculated annual and overall proportions of rural and urban cases per district. Where cases with unspecified localities were included in analyses (Tables 1, 2 and S1), the proportion of unspecified localities were indicated. Annual and overall relative risks (RR) of dengue for each individual district were calculated using:

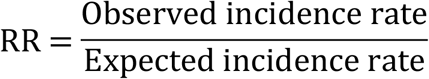

where the expected incidence rate for each district is based on the mean rate for the state multiplied by the population of each district. A RR value > 1 indicates increased incidence of dengue in that location compared to the expected (mean) incidence, and a value < 1 indicates lower than expected dengue incidence.

**Table 1.** Summary of population and dengue burden across Sabah, 2010-2016.

| District | Human population (2010) | Population density (people/km<sup>2</sup>) | Proportion of cases from urban localities* | Mean annual incidence (per 100,000) | Severe dengue mean annual incidence (per 100,000) | Overall relative risk | Number of dengue deaths |
| --- | --- | --- | --- | --- | --- | --- | --- |
| Beaufort | 66,406 | 38 | 0.06 | 49 | 0.2 | 0.9 | 0 |
| Beluran<sup>#</sup> | 106,632 | 14 | 0.14 | 34 | 0.9 | 0.8 | 1 |
| Keningau | 177,735 | 50 | 0.11 | 57 | 0.2 | 0.9 | 0 |
| Kinabatangan | 150,327 | 23 | 0.04 | 19 | 0.0 | 0.3 | 0 |
| Kota Belud | 93,180 | 67 | 0.02 | 52 | 0.7 | 1.4 | 0 |
| Kota Kinabalu | 462,963 | 1,315 | 0.85 | 57 | 0.2 | 1 | 5 |
| Kota Marudu | 68,289 | 36 | 0.02 | 79 | 0.4 | 2.1 | 1 |
| Kuala Penyu | 19,426 | 43 | 0.06 | 161 | 0.0 | 3.5 | 0 |
| Kudat | 85,404 | 66 | 0.50 | 52 | 0.5 | 1.8 | 1 |
| Kunak | 62,851 | 55 | 0.62 | 33 | 1.3 | 1 | 0 |
| Lahad Datu | 206,861 | 28 | 0.42 | 45 | 0.8 | 1.3 | 2 |
| Nabawan<sup>^</sup> | 32,309 | 5 | 0.00 | 88 | 0.0 | 1.6 | 0 |
| Papar | 128,434 | 103 | 0.20 | 31 | 0.0 | 0.6 | 0 |
| Penampang | 125,913 | 270 | 0.51 | 74 | 0.6 | 1.3 | 1 |
| Pitas | 38,764 | 27 | 0.29 | 36 | 0.0 | 0.8 | 0 |
| Putatan<sup>**</sup> | 55,864 | 1,397 | 0.32 | 73 | 0.0 | 1 | 2 |
| Ranau | 95,800 | 26 | 0.09 | 37 | 0.1 | 0.7 | 1 |
| Sandakan | 409,056 | 180 | 0.80 | 57 | 1.0 | 1.1 | 4 |
| Semporna | 137,868 | 120 | 0.53 | 46 | 1.1 | 1 | 2 |
| Sipitang | 35,764 | 13 | 0.14 | 56 | 0.0 | 1.2 | 0 |
| Tambunan | 36,297 | 27 | 0.00 | 37 | 0.0 | 0.8 | 0 |
| Tawau | 412,375 | 67 | 0.73 | 33 | 0.8 | 0.8 | 5 |
| Tenom | 56,597 | 13 | 0.12 | 84 | 0.7 | 1.5 | 0 |
| Tongod<sup>**</sup> | 36,192 | 4 | 0.05 | 22 | 1.0 | 0.4 | 0 |
| Tuaran | 105,435 | 90 | 0.26 | 56 | 0.4 | 1.3 | 0 |
| **Total** | **3,206,742** | **43** | **0.48** | **50** | **0.5** |  | **25** |

\*Rural urban locality data was included between 2011-2016; overall proportions were 0.48 urban, 0.44 rural and 0.08 unspecified.
<sup>#</sup> Beluran was formerly known as Labuk Sugut. <sup>^</sup> Nabawan was formerly known as Pensiangan.
<sup>\*\*</sup> Putatan and Tongod districts only commenced notifications in 2012.

**Table 2.** Mosquito larvae species collected from case residences in Sabah during 2015-2016.

| Locality | Total number of cases in 2015-2016 | No. of case residences inspected | No. of larvae-positive case residences (all species) | House Index (HI) | Number of larvae-positive residences with specific species present (HI) |  |  |  |  |
| --- | --- | --- | --- | --- | --- | --- | --- | --- | --- |
|  |  |  |  |  | *Ae. aegypti* | *Ae. albopictus* | *Ae. aegypti* & *Ae. albopictus* * | *Culex* spp. | Undetermined |
| Urban residences | 3,157 | 255 | 206 | 0.81 | 47 (0.23) | 142 (0.69) | 7 (0.03) | 6 (0.03) | 4 (0.02) |
| Rural residences | 2,824 | 437 | 388 | 0.89 | 33 (0.09) | 225 (0.58) | 4 (0.01) | 26 (0.07) | 100 (0.26) |
| Locality unspecified | 565 | 27 | 24 | 0.89 | 3 (0.13) | 16 (0.67) | 0 (0.0) | 1 (0.04) | 4 (0.17) |
| **Total** | **6,546** | **719** | **618** | **0.86** | **83 (0.13)** | **383 (0.62)** | **11 (0.02)** | **33 (0.05)** | **108 (0.17)** |

HI = proportion of residences positive for mosquito larvae, calculated as number of residences with larvae/number of residences inspected.
\* Both species found breeding together in one household.
Species were undetermined if the larvae failed to survive to adults to be identified, or if identification was pending/incomplete.

## Results

### Temporal trends across the state

A total of 11,882 dengue cases were reported in Sabah during the 7-year study period, with 25 deaths. Cases were reported year-round, with outbreaks commonly occurring in the second half of the year between July and December, sometimes continuing into January and February (Fig 2). Seasonal decomposition analysis showed that, on average, notifications peaked each January, with the highest risk period being between November and March. Smaller peak periods were also observed occasionally in July and October (S1 Fig).

**Fig 2.**
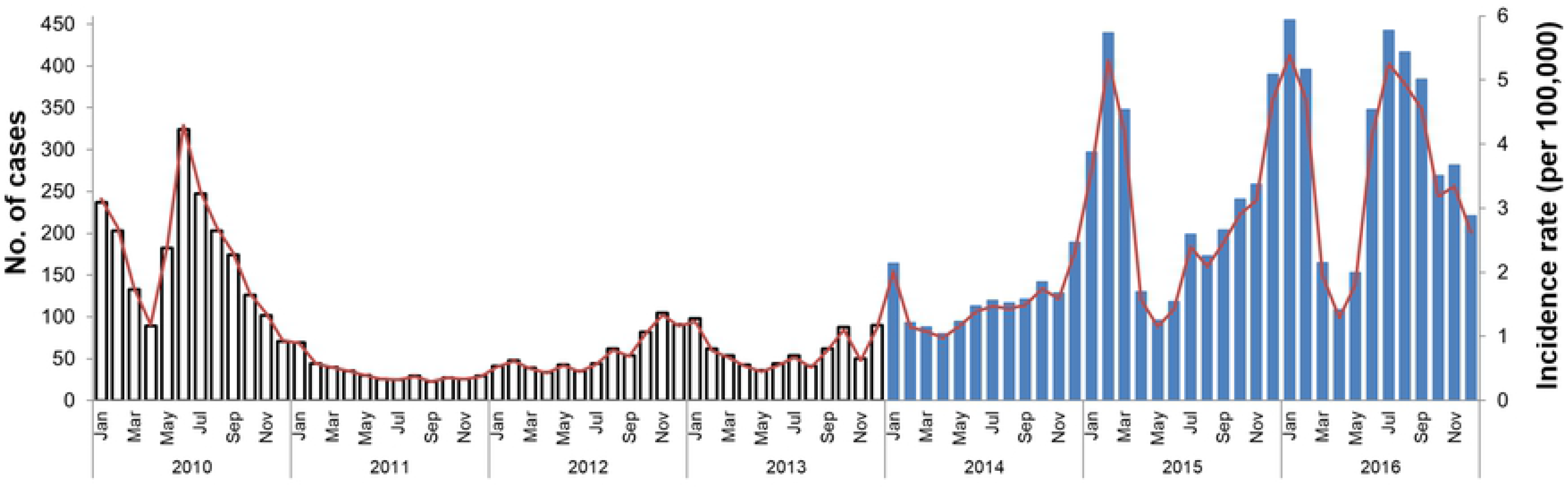
Temporal pattern of dengue in Sabah, 2010-2016. The monthly number of reported dengue cases per year are shown (primary vertical axis), and the corresponding monthly incidence rate (secondary vertical axis). The change in case definition during the study period is indicated by different colour bars: grey bars during the years 2010-2013 where case diagnoses were predominantly clinically-based (with or without laboratory confirmation), and blue bars for the period 2014-2016 where all cases were laboratory confirmed.

Outbreaks varied in magnitude between years, with the largest outbreaks in 2010 and from 2015-2016 (Fig 2). During these large outbreak years, state-wide annual incidence peaked at between 35 and 43 cases per 100,000, respectively. Conversely, incidence rates dropped to 5-9 per 100,000 during the smaller outbreak years between 2011 and 2013. The mean state-wide annual incidence rate across the 7 years was 21 cases per 100,000 people.

For the period 2011-2016, state-wide mean annual incidence of dengue in urban localities was 44/100,000 versus 47/100,000 for rural localities, and annual rates of dengue in urban and rural localities often contributed similarly to the overall burden (Fig 3). However, there was a notable difference during the large outbreaks of 2015 and 2016, when the highest incidence localities appeared to switch between being predominantly urban in 2015 to predominantly rural in 2016.

**Fig 3.**
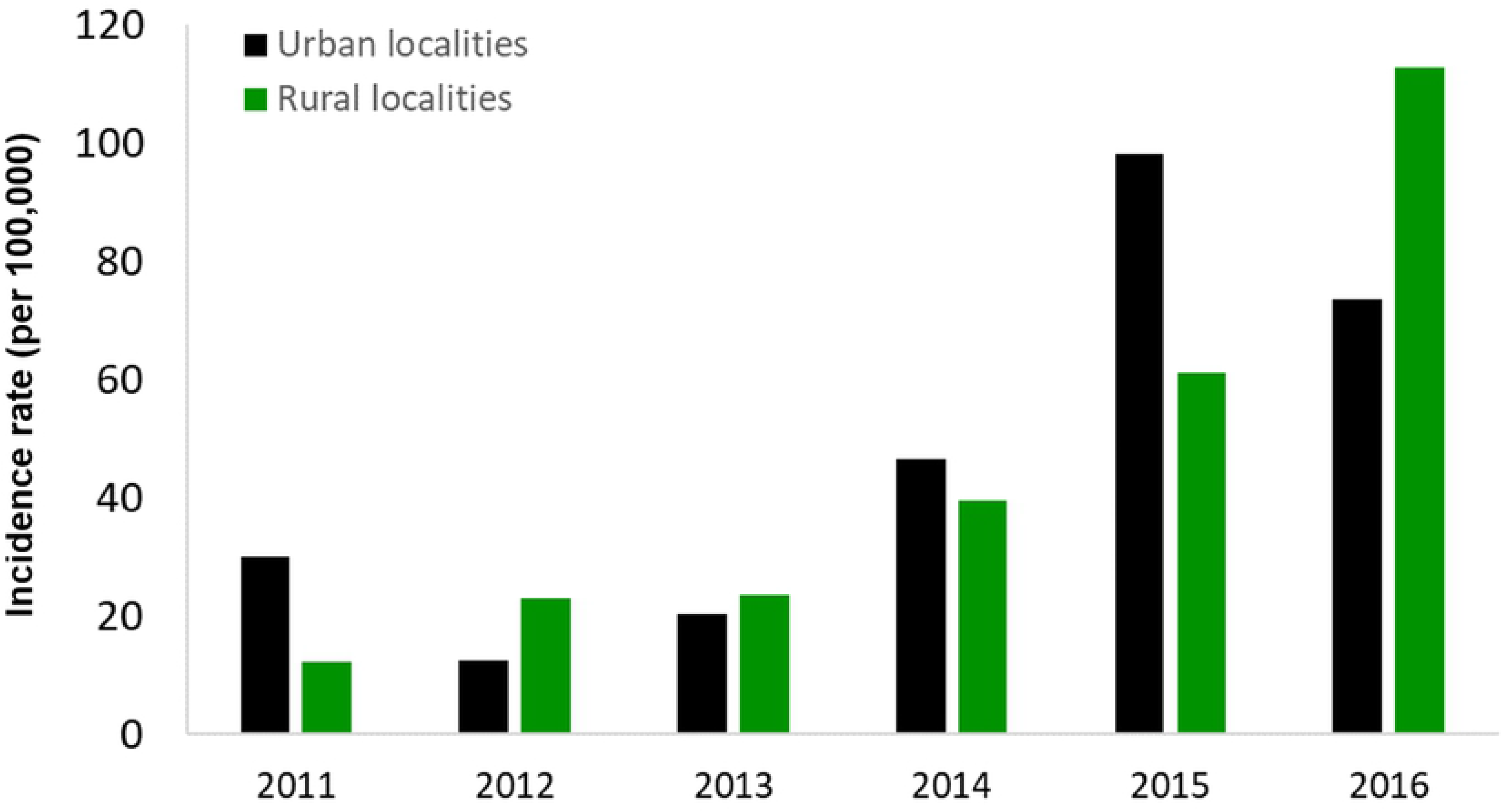
State-wide annual incidence of dengue in rural and urban localities, 2011-2016. Annual incidence rates across the state are shown for cases residing in either urban or rural localities, over a 6-year period.

### Demographic trends

Analyses of demographic trends across Sabah indicated a slightly higher proportion of male dengue cases (60%) than females (40%). After adjusting for differences in population proportions, incidence rates were not significantly different between the two (29/100,000 for males and 20/100,000 for females, p=0.32; Fig 4). This was relatively consistent across all Sabah districts; however, there were some districts with above-average proportions of male cases – in particular, in Tongod and Kinabatangan (75% and 65% male cases, respectively).

**Fig 4.**
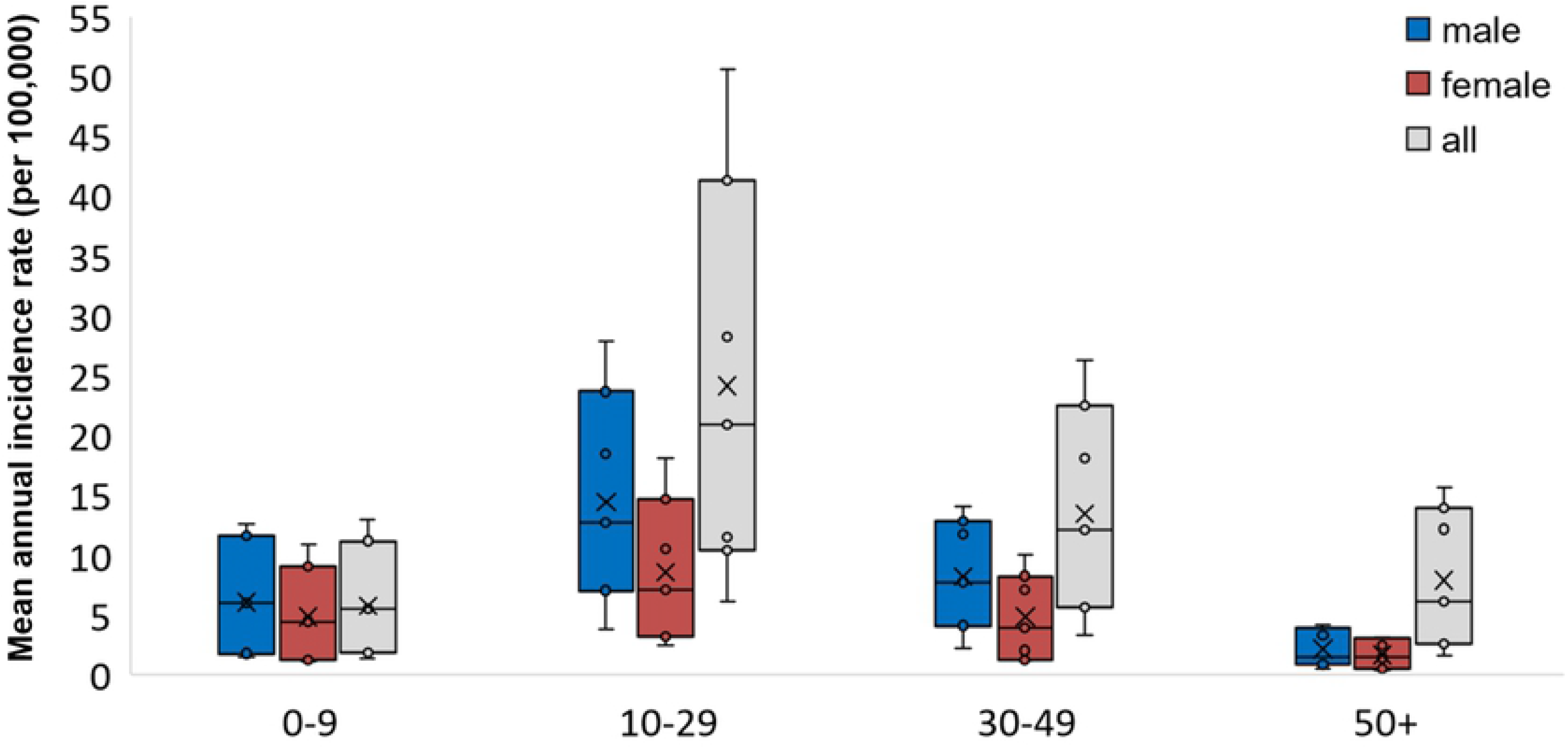
Incidence of dengue in Sabah by age group and gender, 2010-2016. Age- and gender-adjusted incidence rates across Sabah during the 7-year period are shown for males, females and for both genders. * Age-group 10-29 for both genders had a statistically significantly higher mean rate, p=0.029.

Older children and young adults were the dominant age groups affected by dengue (Fig 4), with the majority of cases occurring between 10 and 29 years (mean annual incidence of 24 cases/100,000; 47% of the burden across age groups), followed by 30-49 years (mean 13 cases/100,000/year; 26% of total cases). The median age of all notifications was 25. After adjusting for differences in population proportions, the 10-29 age group had significantly higher incidence than the other age groups (p=0.029). The lowest proportion of notifications occurred below 10 years of age (mean 6/100,000/year), followed by those 50 years of age and above (8/100,000/year).

### Spatial trends across districts

District-level incidence rates were highly variable each year, with a mean annual rate of 50 cases/100,000 (range 19-161 cases/100,000) across the 7 years (Table 1, S2 Fig). High annual variability meant that there was here was no significant difference in mean incidence rates between the districts overall (p=0.462); however, the highest mean incidence rates were found in districts in the west of the state with relatively low human population density, including Kuala Penyu, Nabawan, Tenom and Kota Marudu. These high incidence districts also displayed the greatest extremes in annual dengue rates, ranging between 15 and 942 cases/100,000 each year (S2 Fig). The overall relative risks were highest in Kuala Penyu, Kota Marudu and Kudat districts (RR=3.5, 2.1 and 1.8, respectively; Table 1). The 4 highest-incidence districts reported a low proportion of cases residing in urban localities (0-12%; Table 1). Lower, less variable incidence rates were recorded from some of the central and eastern districts including Kinabatangan, Tongod, Kunak and Tawau (annual incidence range of 3-62 cases/100,000 each year). These districts also had some of the lowest relative risks (RR=0.3, 0.4, 1, and 0.8, respectively; Table 1, S2 Fig), along with a wide range in their proportions of urban cases (4-73%).

The changing annual spatial trend is shown in Fig 5, which shows high annual and mean incidence rates often occurring in the western districts of Sabah. A shift in dynamics occurred during the large 2015 outbreak, when incidence increased markedly in the more densely populated western districts of Kota Kinabalu, Penampan, Putatan, and in Sandakan and Semporna in the east (Fig 5). The overall urban case proportions in these districts ranged from 53-85%. During 2016, cases from both urban and rural localities contributed to the outbreak, but the greatest overall incidence was in Tenom, Nabawan and Keningau districts, where the majority of cases were from rural localities.

**Fig 5.**
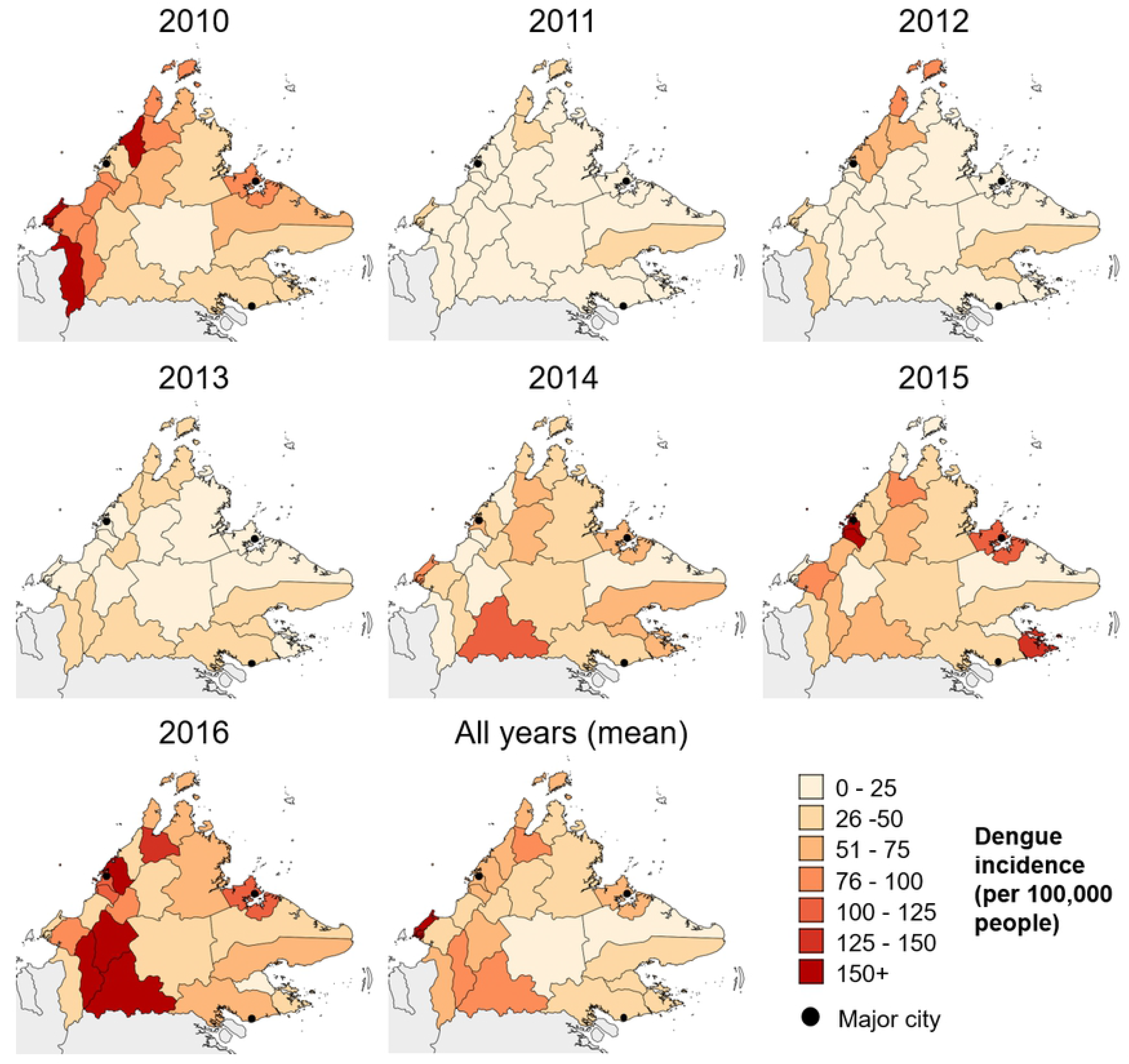
Annual spatial incidence of dengue in Sabah, 2010-2016. Incidence rates across districts are shown for each year, as well as the overall mean annual incidence during the 7-year period. The 3 major cities of Sabah (Kota Kinabalu, Sandakan and Tawau) are indicated by black circles.

### Severe dengue

Of all dengue cases reported over the 7 years, 1.1% were severe (haemorrhagic) dengue cases. The average annual state-wide number of severe cases was 18, although this increased to 28 and 25 cases during 2011 and 2013, respectively, despite these being relatively low incidence years (Figs 2, 6). The greatest proportion of severe cases were concentrated in Sandakan (24%) and Tawau (20%) districts on the eastern side of the state, with the highest severe dengue incidence found in Kunak, Sandakan and Tongod (Fig 6; Table 1). The lowest proportion and incidence of severe dengue was observed in the western districts, several of which recorded zero severe cases, despite recording high overall dengue incidence (Figs 5, 6; Table 1). Severe dengue occurred evenly across both genders and age groups, although the burden was highest for age groups under 30 years, with the largest proportion (35%) reported within the 10-19 years age group. There were 9 severe dengue deaths during the study period, 4 of which were in Tawau district. Deaths from severe dengue occurred consistently across years, genders and age groups.

**Fig 6.**
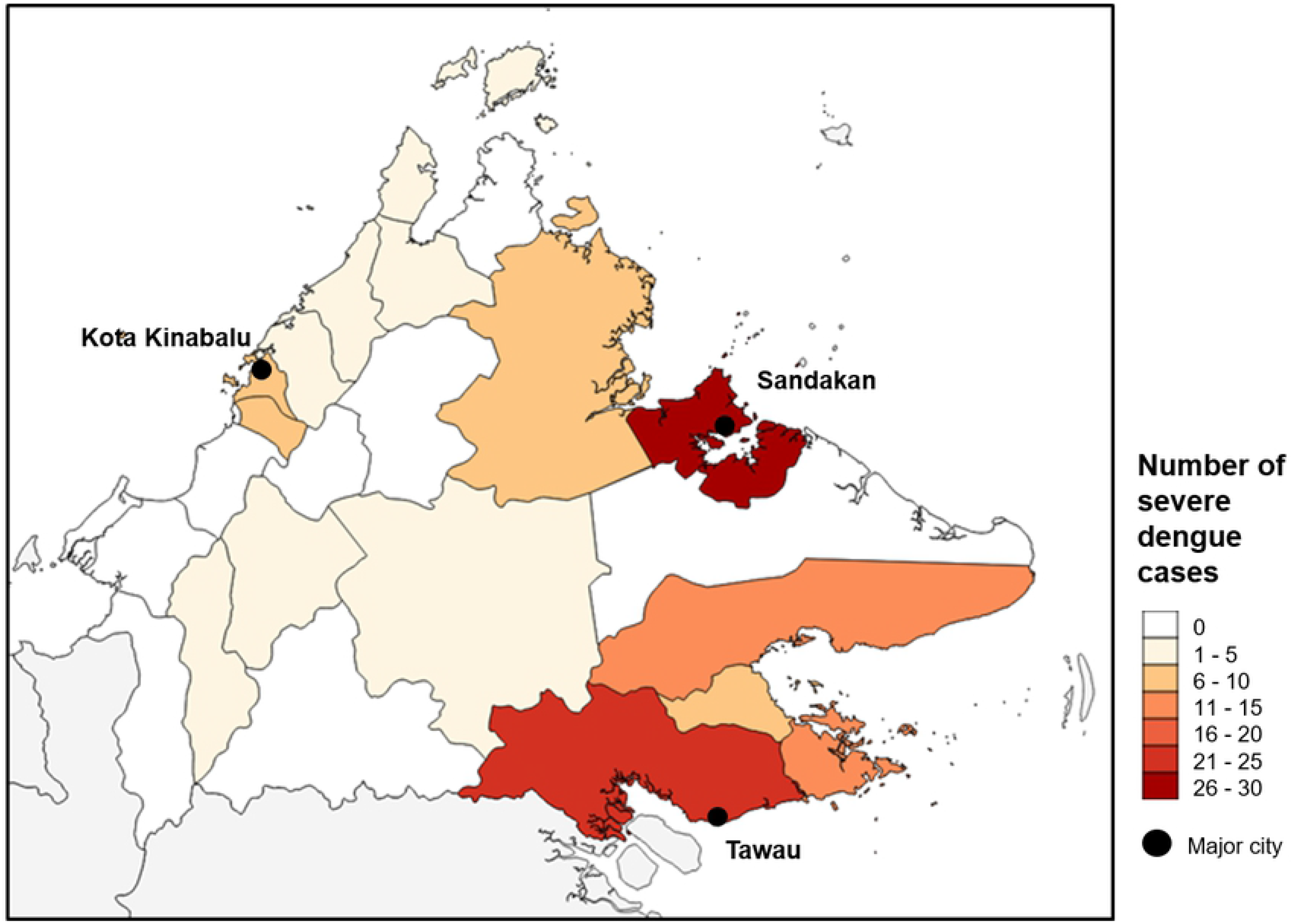
Total severe dengue notifications by district, 2010-2016. Total number of severe dengue cases reported for each district during the 7-year period. The 3 major cities of Sabah (Kota Kinabalu, Sandakan and Tawau) are indicated.

### Entomological factors

Entomological data collected within the 2015-2016 outbreaks indicated that the partially sylvatic vector *Aedes albopictus* was the predominant species detected in larval collections from both rural and urban case residences (Table 2). Of 719 dengue case residences that were inspected as part of active surveillance in 2015-2016, 618 were found to contain mosquito larvae (HI=86%). Of those, *Ae. albopictus* larvae were identified from 394 residences (HI=64%), either alone (383 residences) or with *Ae. aegypti* (11 residences). Conversely, 94 residences were positive for *Ae. aegypti* (83 alone, 11 with *Ae. Albopictus;* HI=15%), 33 were positive for *Culex* species (HI=5%), and 108 larval samples could not be identified (17%).

The specific districts where case residences were inspected are detailed in S1 Table. Inspections were conducted in 21/25 districts, though the majority were conducted in Tawau (158 inspections; with 96 in urban localities) and Nabawan (107 inspections; all in rural localities). The majority of residences positive for *Ae. aegypti* were in the east coast districts of Tawau (larvae found in 60/158 residences) and Lahad Datu (18/43 larvae-positive residences). Residences in Tawau and Lahad Datu districts together comprised 80% of all Sabah residences where *Ae. aegypti* was identified. *Ae. albopictus* was prevalent across both urban and rural residences of most of districts surveyed. The majority of residences positive for *Ae. albopictus* larvae were also in Tawau (101/158 residences), followed by Penampang (52/58 larvae-positive residences) and Nabawan and Keningau (46 residences each; S1 Table).

## Discussion

### Spatial and temporal trends

In recent years, the scale of epidemics in South East Asia has increased in both urban and rural areas (28–32). We found an overall increasing incidence trend in the Malaysian state of Sabah, with the highest risk period occurring annually between November and March. While spatial trends in dengue incidence varied from year to year, the most intense transmission across all years occurred in districts along the western coast of Sabah. The timing of large epidemic years in Sabah (2010, 2015 and 2016) was consistent with patterns observed at national and regional levels during the same period (wider Malaysia, Indonesia, Philippines) (33, 34). This suggests shared seasonal influences on outbreak occurrence although, especially in the tropics, the causative relationship between temperature, rainfall and transmission remains poorly understood (35, 36).

We also noted consistently high incidence in rural localities across the study period, as well as a shift in the spatial dominance of urban versus rural localities during the large outbreaks of 2015-2016. Our findings suggest that high density, urbanised areas are not necessarily the primary drivers of ongoing epidemics in Sabah, and that factors other than population size may drive the risk in rural areas. Recent studies in other areas of Malaysia have also shown that dengue infection can occur at equivalent rates in both rural and urban areas (37, 38). It could be that the threshold human density required to maintain transmission may be lower than previously thought, although human movement (which may relate to population density) between urban and rural areas is also likely to have influenced the patterns we observed (39, 40).

Rural dominance of dengue has also been observed elsewhere in the region, including in Cambodia, Thailand, Vietnam and Sri Lanka (41–44). These countries have all reported epidemics spreading between rural and urban areas – in both directions – via human or mosquito movement, facilitated by favourable climatic conditions. Other potential influences on both urban and rural dengue transmission in Sabah might include water storage practices, mosquito vector ecology and sociocultural factors (32, 45–48). The relative importance of some of these risk factors in mediating dengue transmission is still not well understood, even in urban areas (49, 50). Understanding the relationships between these risk factors may be challenging to disentangle in different environments, and especially where urban and rural areas are highly interconnected; however, knowledge of these dynamics may be important to optimise the design and targeting of dengue control strategies. This is especially important given the potential for ongoing outbreaks in both urban and rural areas of Sabah.

### Demographic factors

Our findings indicated that the age-related dengue risk in Sabah was in line with regional trends indicating a transition from children to adults being disproportionately affected by dengue (51). Incidence was higher for males than for females across all districts of the state, and was significantly higher for both genders in the 10-29 age group. This higher risk may suggest that a larger proportion of people in this age-group (and possibly males in particular) were either engaged in outdoor activities and/or being occupationally exposed. The agriculture sector is the major employment sector in Sabah, and this type of work may increase exposure to mosquitoes (52, 53). The particularly high proportion of males affected in Tongod and Kinabatangan districts may reflect the fact that both are very rural, and may have a larger proportion of men engaged in agricultural or recreational outdoor activities. Outdoor activities, especially those in close proximity to forests or forest edges, are thought to increase the risk of being bitten by the abundant exophilic vector, *Ae. albopictus* (54, 55). However, studies in peninsula Malaysia have shown that *Ae. albopictus* can also adapt to indoor urban environments (56, 57); hence, the high proportion of cases we observed in 10-29 year olds could also suggest infection in indoor environments at home or at school. Further investigation to identify the specific factors associated with infection risk in Sabah may be useful to inform prevention strategies for this high-risk group.

### Severe dengue

Changing demographic or immunological factors may also explain the observed pattern of severe dengue in our study. Severe dengue showed a decreasing trend over time (despite the overall increasing trend in incidence rates), but with the highest risk localised to two main regions of the state: the eastern districts of Tawau and Sandakan. These districts include major urbanised cities as well as rural surrounding areas, and comprised relatively low dengue rates compared to the west of the state. The reasons for this spatial concentration of cases in these eastern districts is unknown, though it’s possible that a serotype switch from DENV 4 to DENV 1 reported to have occurred in Sandakan between 2013 and 2016 may have contributed (12). It might also be possible that different areas of Sabah experience serotype changes more or less frequently depending on levels and direction of population movement (33, 58). Surveillance information regarding which virus serotypes and genotypes were circulating in Sabah was not available in this study, so we were unable to assess the potential contribution of virus circulation patterns to the trends we observed. Assessing serological surveillance data alongside epidemiological data in future studies in Sabah could aid predictions of severe disease risk (59, 60).

### Entomological factors

During the large outbreak period between 2015 and 2016, our entomological surveillance data indicated a striking association between the presence of the mosquito vector *Ae. albopictus* relative to *Ae. aegypti*, in both urban and rural case residences in the majority of the state. Interestingly, the eastern districts of Sabah state appeared to have a higher proportion of *Ae. aegypti* compared to the rest of the state, although overall dengue incidence was lower on the east coast. Interestingly, this finding was consistent with those of early entomological surveys of Sabah in the 1970’s, which reported higher numbers of *Ae. aegypti* on the east coast and lower abundance on the west coast (61, 62). In those studies, the greater presence of *Ae. aegypti* in the east was thought to be due to more frequent travel by boat between east coast settlements for fishing and trade.

Although *Ae. aegypti* is generally considered responsible for most dengue transmission in South East Asia (63, 64), *Ae. albopictus* is more common than *Ae. aegypti* across Malaysia and Borneo (52, 54, 55, 65). Its competence for specific dengue genotypes, its abundance in both rural and urban areas, its biting behaviour and its diverse aquatic habitat may all account for patterns of mosquito-human contact and subsequent transmission in Sabah (48, 66). The presence of natural and artificial larval habitats for *Ae. albopictus* have previously been associated with epidemic disease in both urban and rural areas of Malaysia (56, 57, 67) despite the fact that globally, *Ae. aegypti* is undoubtedly the predominant vector driving epidemics (68, 69). Urban dominance of *Ae. albopictus* has also been observed, at least seasonally, in parts of Thailand, southern China and other South East Asian countries (70–72). Given the likely role of *Ae. albopictus* in mediating dengue epidemics in Sabah, vector control strategies may have to be expanded to include both *Ae. aegypti* and *Ae. albopictus*. Because *Ae. albopictus* is commonly characterized as more exophagic and exophilic than *Ae. aegypti*, and exploiting a wider range of hosts and habitats in peri-urban and rural environments, targeting outdoor resting sites of adult *Ae. albopictus* may be a useful control strategy in Sabah (73–75).

### Limitations

The main caveat to our findings is that the changes in dengue case definition in Malaysia in 2014 may have influenced the trends reported here, in terms of either under- or over-reporting of cases. Reduced reliance on clinical symptoms for case notification from 2014 onwards would be expected to reduce notifications dramatically but, in fact, a dramatic increase in cases were recorded. It is possible that prior to that date, the lack of resources for testing or notifying dengue, or other socioeconomic factors, may have resulted in under-reporting (76). It is also possible that increases in diagnostic testing from 2014 were not uniform across all districts, and/or that additional reporting inconsistencies may have impacted our observations.

### Conclusions

The rising magnitude of dengue in Sabah in both rural and urban areas suggests that a better understanding of dengue transmission across different environments is needed. Our findings support the notion that dengue epidemics can be both urban and rural environment-driven, and suggest risk factors that may be of use for clinicians, public health practitioners and vector control teams. The trends observed in Sabah indicate that localized ecological, human, virus and vector dynamics may be predictive of dengue epidemics irrespective of urban and rural environment. In Sabah, as with many countries of South East Asia, there is likely a complex interplay of these factors operating in both rural and urban areas, and these probably overlap (10, 29). Considering the ongoing expansion of dengue endemicity and burden in the region, proactive strategies to increase understanding of the complex and evolving epidemiological factors underlying dengue risk across varied environments are critical.

## Acknowledgements

The authors would like to acknowledge the contribution of the Sabah Department of Health, Ministry of Health, Malaysia for making dengue notification data available. We also thank the Director General of Health Malaysia for the permission to publish this paper. We are grateful for assistance and input provided by Nicholas Anstey, Matthew Grigg, Kimberley Fornace, Christopher Wilkes, Eloise Stephenson and Andrea Rabellino. The author(s) received no specific funding for this work. The authors declare no conflict of interest in conducting this study.

## Supporting Information

**S1 Fig.**
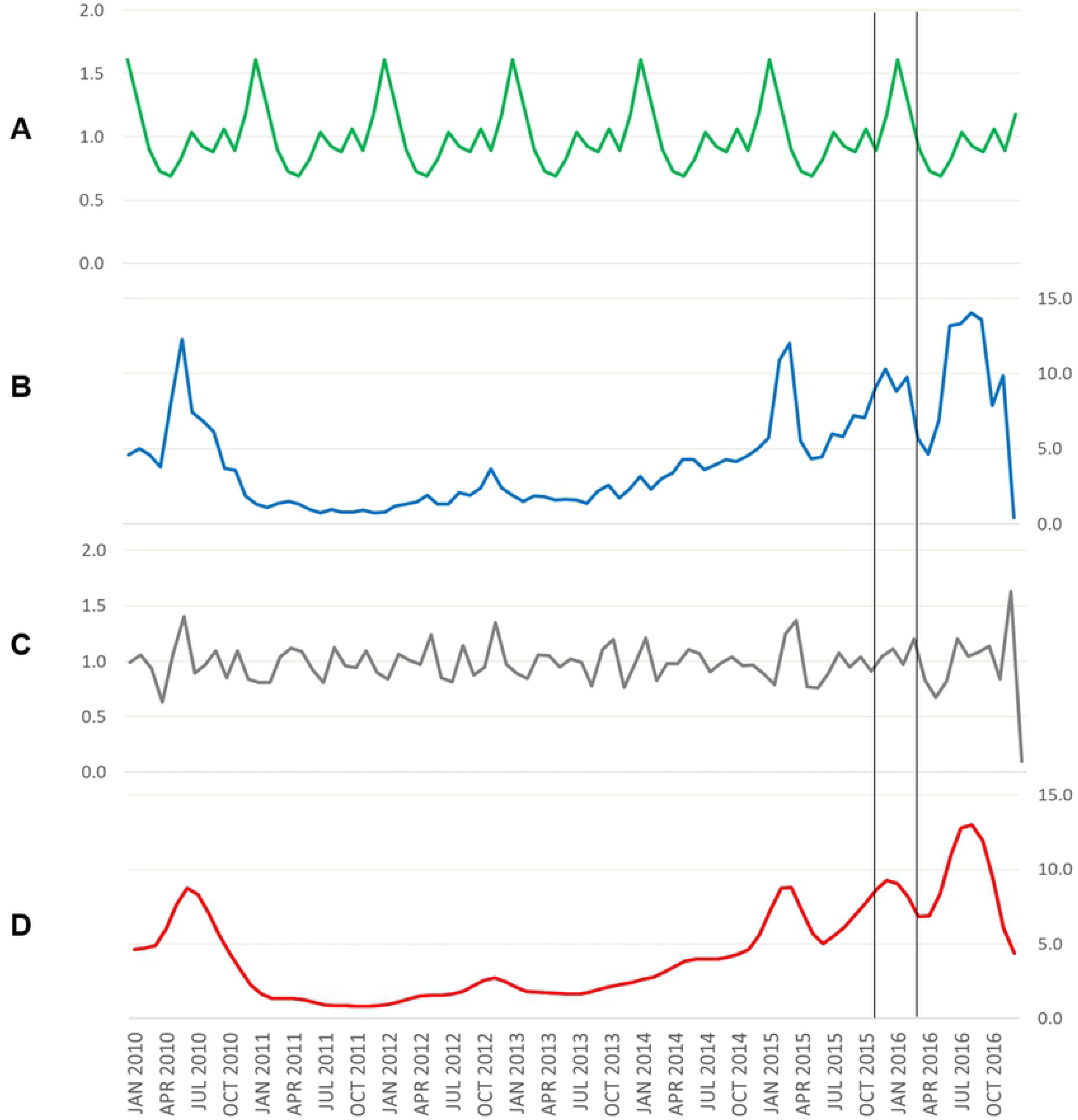
Seasonal decomposition of incidence rates in Sabah, 2010-2016. The seasonal trend of dengue is shown in panel A, with the largest seasonal peak occurring on average between Nov and May each year (indicated by vertical black lines). The additional components separated from the seasonal trend during the decomposition procedure are also indicated in panels B-D (cyclical component (B), irregular component (C) and overall smoothed trend (D)).

**S2 Fig.**
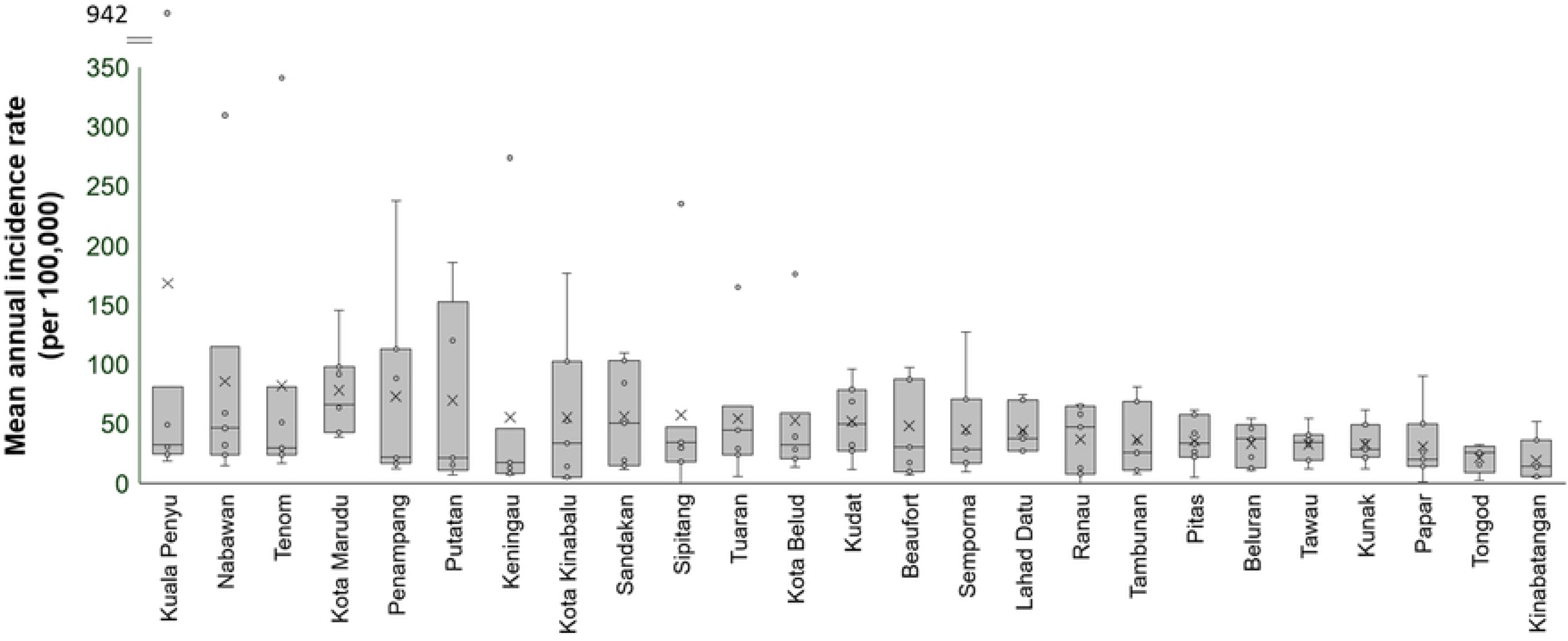
Variation in dengue incidence across Sabah districts, 2010-2016. Dengue mean annual incidence rates across all years are plotted, in order of highest-lowest mean annual incidence rate, showing the mean (x), median (line), and the range of rates (upper and lower whiskers) across the years.

**S1 Table. Entomological surveillance of case residences by district, 2015-2016.**

**S1 Checklist: STROBE Checklist for observational studies.**

